# ABC transporters confer multidrug resistance to Drosophila intestinal stem cells

**DOI:** 10.1101/511584

**Authors:** Hannah Dayton, Jonathan DiRusso, Kristopher Kolbert, Olivia Williamson, Aiste Balciunaite, Edridge D’Souza, Kelly Becker, Elizaveta Hosage, Muneera Issa, Victoria Liu, Raghuvir Viswanatha, Shu Kondo, Michele Markstein

## Abstract

Adult stem cells can survive a wide variety of insults from ionizing radiation to toxic chemicals^1–3^. To date, the multidrug resistant features of stem cells have been characterized only in vertebrates, where there is a critical need to understand how cancer stem cells thwart chemotherapy drugs^4–6^. These studies reveal that the ability of both normal and cancer stem cells to survive toxins hinges on their high levels of expression of ABC transporters, transmembrane pumps that efflux lipophilic compounds out of cells^7,8^. This has been observed across a wide spectrum of vertebrate stem cells including breast, blood, intestine, liver, and skin, suggesting that high efflux ability and multidrug resistance may be general features of stem cells that distinguish them from their differentiated daughter cells. Here we show that these previously described vertebrate stem cell features are conserved in Drosophila intestinal stem cells. Using a novel in vivo efflux assay and multiple drug challenges, we show that stem cells in the fly intestine depend on two ABC transporters—one constitutively expressed and the other induced—for efflux and multidrug resistance. These results suggest that stem cell multidrug resistance by ABC transporters is a general stem cell feature conserved over 500 million years of evolution.

---

All cells from bacterial cells to human cells express ABC transporters^9^. Vertebrate stem cells and their cancer stem cell counterparts, however, tend to express ABC transporters at unusually high levels, endowing them with resistance to a wide array of toxic compounds including most chemotherapy drugs^4–6^. Currently it is not known how ABC transporter gene expression is regulated in stem vs. daughter cells, nor whether there are costs associated with their expression, as recently found in bacterial cells^10^, that limit their levels and cell types of expression. These questions can be answered ideally in simple model organisms. However, to date stem cell multidrug resistance has been described only in vertebrates. Here, we show that that the property of multidrug resistance extends to Drosophila intestinal stem cells. This finding points to the generality of stem cell multidrug resistance from vertebrates to invertebrates and establishes a new model that can be fully dissected with the power of Drosophila genetics.

To identify Drosophila stem cells with drug resistant properties, we focused on the stem cells that maintain the adult fly intestine, as these cells are optimally positioned to come in contact with ingested compounds. Like our intestine, the fly intestine is composed of a single layered epithelial tube that is regionalized along its length for distinct digestive and absorptive functions^11,12^. We focused on the R5 region, which is analogous to the mammalian small intestine and specialized for absorption^13^ (Figure 1a). The major differentiated cell type in this region, large polyploid enterocytes (ECs), form a brush border epithelium at their apical ends^14^ which, like in mammals^15^, is specialized for carrying out absorptive functions (Figure 1a). When ECs die, intestinal stem cells (ISCs)^14,16^ in the epithelium divide, resulting in two daughter cells: one maintains its stem cell identity and the other, called an enteroblast (EB) or progenitor, matures into an EC^17–20^ (Figure 1b). The ISC and progenitor cells can be distinguished by their expression of the transcription factor escargot^14^, which has been shown to be necessary and sufficient the maintain the stem cell state(MIR8PAPERetc). The ECs can be distinguished by their expression of the brush border cytoskeletal protein Myosin 1A^21^. In the present work, we exploit Gal4 enhancer traps in escargot (esg-Gal4)^14^, and Myosin 1A (Myo1a-Gal4)^17^ to mark and manipulate the stem/progenitor cells and ECs respectively (Figure 1b).

**Figure 1.**
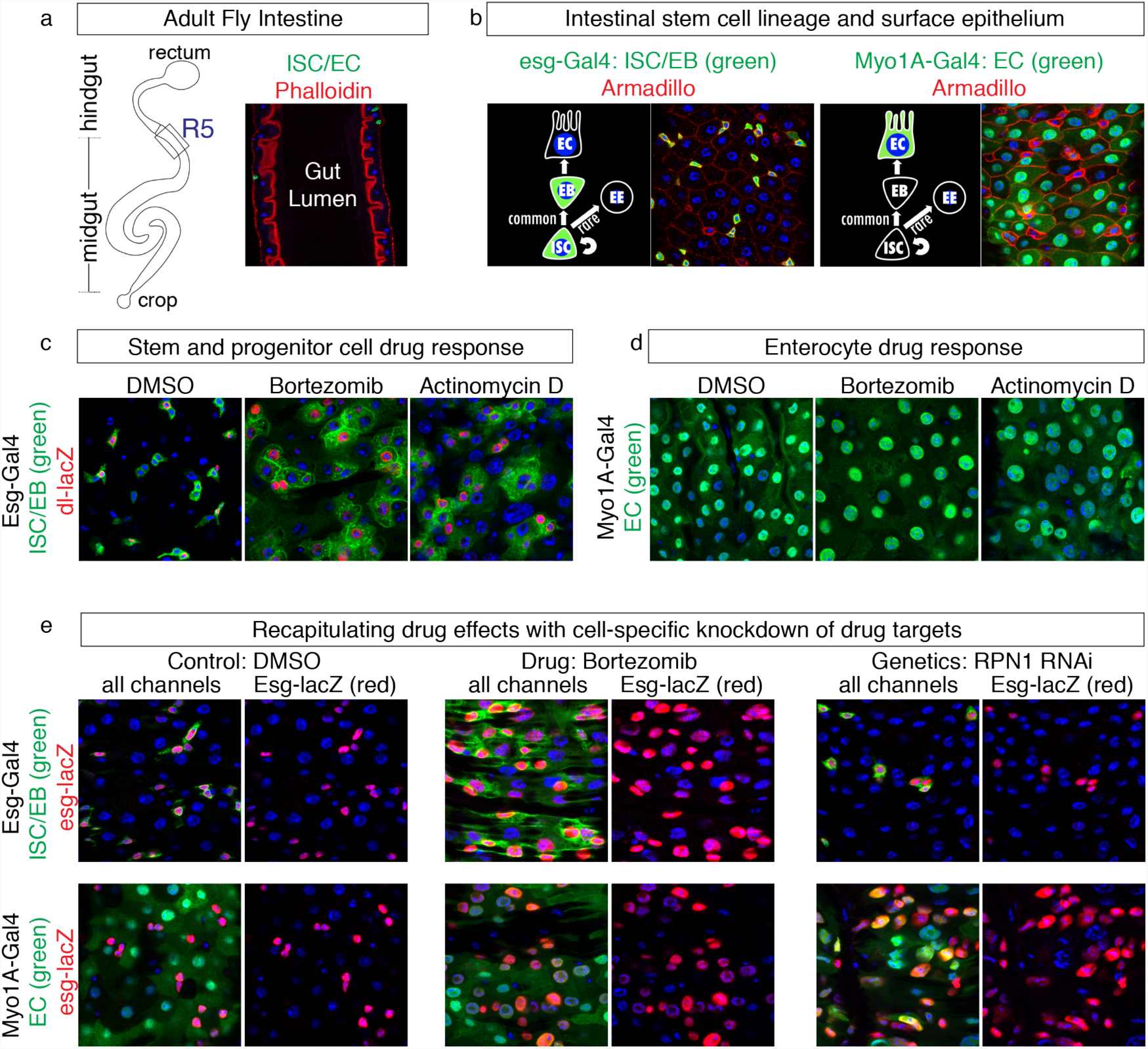
Evidence that drugs do not reach their targets in intestinal stem progenitor cells. **(a)** Schematic and confocal cross-section showing the R5 region of the adult Drosophila intestine. Phalloidin staining of F-actin shows rich brush border of ECs lining the gut lumen and muscle lining the outside of the gut. Stem/progenitor cells marked in green are positioned basally abutting the surrounding muscle. Nuclei in this image and throughout the paper are stained with DAPI (blue). **(b)** Intestinal stem cell lineage and surface views of the gut epithelium showing stem and progenitor cells marked by esg-Gal4 driving GFP and ECs marked by Myo1A-Gal4 driving GFP. **(c)** Effect of drugs on ISC (dl-lacZ, red) and progenitor (not red) esg+ green cells. **(d)** Effect of drugs on ECs (green). **(e)** Effect of the proteasome targeting drug bortezomib (middle panels) and RNAi against the proteasome subunit RPN1 driven by esg-Gal4 (top right panel) and by Myo1A-Gal4 (bottom right panel). Esg+ cells marked with esg-lacZ in red in all e-panels.

To investigate the differential effect of cytotoxic drugs on ISCs vs. ECs, we used the stem cell specific reporter dl-lacZ^17,22,23^ to track stem cells in flies treated with two structurally and physiologically distinct chemotherapy drugs, bortezomib, a proteasome inhibitor, and actinomycin D, a transcription inhibitor. In previous work, we had shown that these drugs induce ECs to express the Jak-Stat cytokine Upd3 which in turn induces esg+ cells to proliferate^24^. However, we did not distinguish the stem cells from progenitors nor assess the effects on EC survival. As shown in Figure 1, dl-lacZ+ ISCs increase in number showing that that they are able to not simply survive, but “thrive” in the presence of cytotoxic agents (Figure 1c). The increase in esg+ dl-lacZ-cells indicates that progenitor cells also increase number in the presence of cytotoxic drugs. Conversely, we found that these same drugs killed about 50% of the ECs. The remaining ECs exhibited increased ploidy and size, consistent with previous observations showing that genetically induced EC death results in expansion in the size of remaining ECs^25^. These observations suggest that cytotoxic drugs reach their targets in ECs but not stem or progenitor cells.

To test whether cytotoxic drugs in fact reach their targets in stem and progenitor cells, we took a genetic approach using the Gal4-UAS TARGET system^26,27^. Specifically, we expressed RNAi against known drug targets in stem and progenitor cells with esg-Gal4 to recapitulate the effect of drugs reaching their targets in these cells. We also expressed RNAi against drug targets in the ECs with Myo1A-Gal4. As shown in Figure 1, RNAi knockdown of RPN1^28^, a proteasome subunit, recapitulates the effect of feeding flies the proteasome inhibitor bortezomib, when the RNAi is expressed in ECs, but not when it is expressed in stem and progenitor cells (Figure 1e). In fact, RPN1 expression in stem and progenitor cells results in about a 50% decrease in esg+ cells (marked with esg-lacZ), indicating that bortezomib, which causes an increase in esg+ cells, does not reach its target in these cells. We obtained similar results knocking down tubulin84 (tub84), the binding target of several mitotic inhibitors that induce esg+ compensatory growth including vinblastine, vincristine, and paclitaxel^29^ (Supplement Figure 1). These genetic results indicate that cytotoxic drugs reach their targets in ECs but not stem and progenitor cells.

The finding that multiple drugs appear to not reach their targets in esg+ stem and progenitor cells suggests that there must be a general mechanism that protects stem cells from exogenous toxic compounds. One possibility, previously proposed in the literature^18^ is that the EC daughter cells protect stem and progenitor cells from the contents of the gut lumen (see Figure 1a). The protective barrier function of enterocytes in mammals^30^ is well documented and further supported in Drosophila by recent characterization of adherins junctions between ECs^31^. The other possibility, not mutually exclusive with the first, is that the stem cells have intrinsic multidrug resistance properties, as described by the efflux properties of vertebrate stem cells^32^. To examine both possibilities further, we fed flies a library of 10 red fluorescent dyes, nine of which are membrane permeable and are used as indicators of drug efflux^33,34^. These include the rhodamine derivatives Tetramethylrhodamine (TMRM) and Rhodamine 123 as well as seven Syto Red dyes that range in size from 450 to 550 MW. We also included a large membrane impermeable probe, 40kD dextran beads coupled with TMRM.

The dyes localized in three patterns that collectively argue against the EC barrier model of “passive” stem cell multidrug resistance and for the model of intrinsic stem cell multidrug resistance (Figure 2a). For example, Syto Dye 60 and the membrane impermeable 40kD dextran-TMRM beads localized equally well in the esg+ stem/progenitor cells and the EC daughter cells (pattern I). This finding indicates that the ECs do not form an impenetrable barrier. Further studies will be required to discern if compounds pass from the gut lumen *through* or *between* the ECs to reach the esg+ stem/progenitor cells. The remaining 8 dyes showed exclusion from the esg+ stem/progenitor cells which in combination with pattern I suggest that the esg+ stem and progenitor cells can efflux several substrates. Five of these dyes appear to be excluded specifically from the esg+ cells (pattern II) while three appear to be excluded not just by esg+ cells but also their closest neighboring EC daughter cells (pattern III). The proximity of the dye-excluding ECs to esg+ cells suggests that the youngest ECs may retain some dye-exclusion properties of the esg+ stem/progenitor cells. In support of this possibility, we found that young marked clones exhibited the dye exclusion phenotype whereas older clones showed mixed patterns (Figure 2a; Supplement Figure 2).

**Figure 2.**
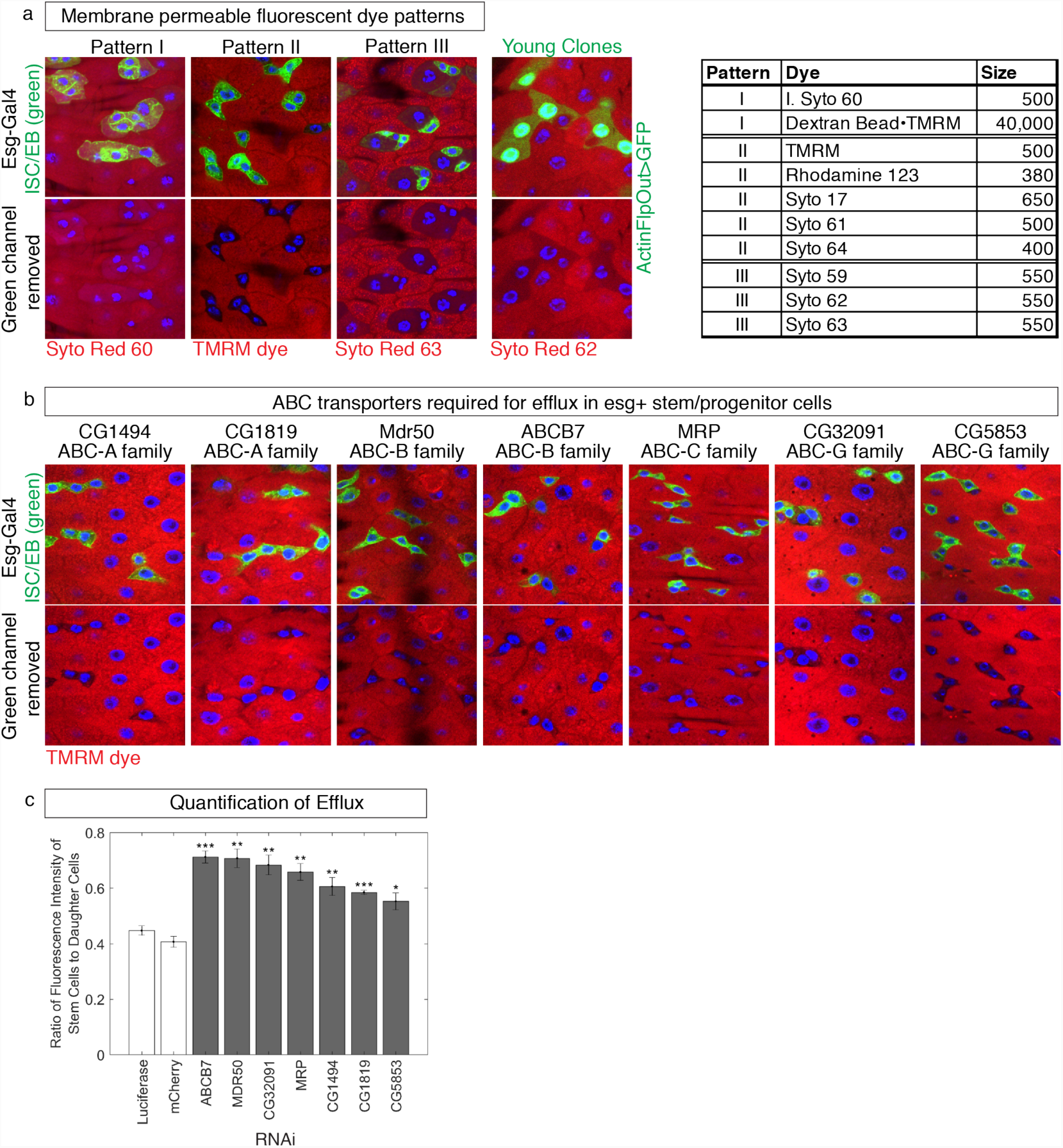
Detection and quantification of efflux by ABC transporters. **(a)** Dye localization observed after two days continuous feeding of red dyes. Stem/progenitor cells marked in green, top panels; green channel removed in lower panels to visualize presence/absence of dye within the stem/progenitor cells. **(b)** Effect of RNAi knockdown on TMRM dye efflux in stem/progenitor cells. **(c)** Ratio of average red fluorescence intensity of per pixel in esg+ stem cell/progenitor cells compared to daughter cells in the same gut. Five or more guts were analyzed per genotype. The error bars represent one standard deviation from the mean. p-values calculated using a 2-sample unequal variance t-test in Excel: *** = p< 0.001, ** = p < 0.01, * = p<0.05.

To test if the esg+ dye exclusion pattern was indicative of efflux by ABC transporters, as widely observed with membrane permeable dye assays in vertebrate stem cells (e.g. see the classic “side-population” assay^35^) we performed an RNAi screen against all 55 ABC transporters encoded in the Drosophila genome, and assessed the effect on efflux using TMRM as a probe (Figure 2a). This screen identified 7 ABC transporters that when knocked down lead to a failure of esg+ stem/progenitor cells to exclude TMRM as seen in pattern II (Figure 2a,b; Supplement Table I). To quantify the effect of each transporter, we measured the ratio of fluorescence intensity per pixel in esg+ cells vs. ECs and found that the knockdown effects were highly reproducible for each transporter, resulting in close to a 2-fold increase in stem/progenitor dye retention (Figure 2e). These results demonstrate that Drosophila stem and progenitor cells, like vertebrate stem cells, can be differentiated from their daughter cells based on possessing greater efflux ability by virtue of ABC transporter activity.

Our finding that seven ABC transporters play essential roles in distinguishing esg+ stem/progenitor cells from ECs, suggests that they may be expressed exclusively in esg+ cells, or simply at higher levels in esg+ cells than in ECs. To distinguish between these possibilities, we characterized their expression patterns by knocking in a T2A-Gal4 reporter^36^ immediately upstream of the stop codon in 5 of the identified transporters (Figure 3a) and we obtained a GFP-mimic enhancer trap^37^ for one of the others (Figure 3b). Of the 6 tagged transporters, we found that two are expressed specifically in stem/progenitor cells: CG32091 and ABCB7 (Figure 3c). CG32091 is expressed in a pattern indistinguishable from the esg pattern under both normal and cytotoxic conditions (Figure 3c). In contrast, ABCB7 expression is not constitutive in the gut epithelium but is instead induced in esg+ stem/progenitor cells by ingested cytotoxins, such as bortezomib and actinomycin D (Figure 3c). Using RTqPCR, we determined that both drugs induce approximately a 3-fold increase in ABCB7 expression (Supplement Figure 3).

**Figure 3.**
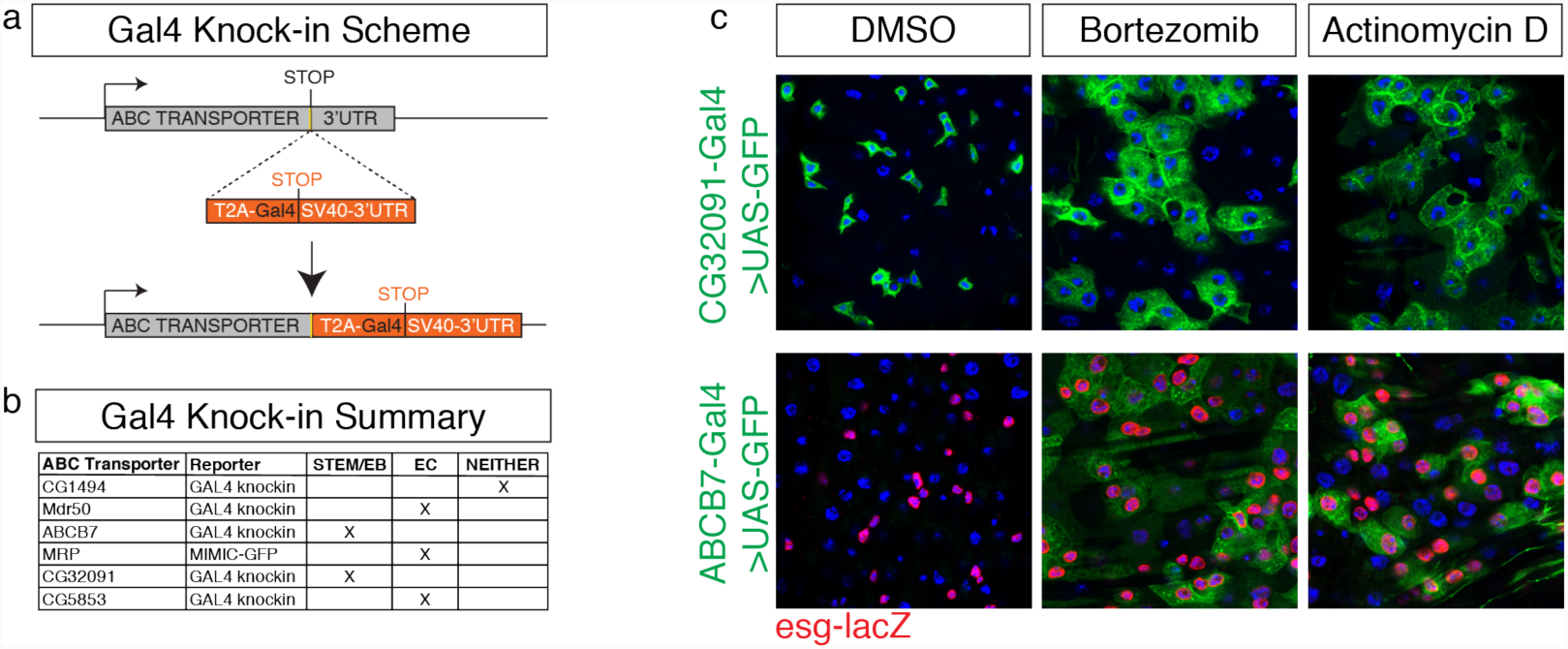
Identification of stem/progenitor specific ABC transporters. **(a)** Scheme for creating T2A-Gal4 knockins immediately after the penultimate codon (yellow) of ABC transporters. This replaces the endogenous 3’UTR with the SV40-3’UTR, creating transcriptional reporters for each gene. **(b)** Summary of expression patterns of ABC transporters identified in the RNAi screen as required for TMRM dye exclusion. Note, some are not visually detectible in the stem/EB cells, yet are required in these cell for dye exclusion, including ABCB7 under normal conditions. **(c)** Expression patterns of CG32091 and ABCB7 under normal (DMSO) and cytotoxic conditions with bortezomib and actinomycin D.

Our finding that ABCB7 and CG32091 are required for dye efflux, and that ABCB7 expression is induced by cytotoxic drugs suggests that one or both might also be required for stem cell multidrug resistance. We therefore tested the ability of esg+ cells to survive and undergo compensatory proliferation in the presence of cytotoxic drugs when each transporter was knocked down by RNAi. As shown in Figure 4, both transporters are required for esg+ cells to undergo compensatory proliferation (Figure 4a) in the presence of cytotoxins. Moreover, knockdown of each transporter results in a 50% reduction in the number of surviving esg+ cells, reminiscent of the effects we observed of knocking down drug targets in esg+ stem/progenitor cells by RNAi (Figure 1e, Supplement Figure 1). In contrast to the results with cytotoxins, ABCB7 RNAi in esg+ cells did not interfere with their ability to undergo compensatory proliferation when challenged with an agent that is not effluxed by transporters, the chemical irritant dextran sodium sulfate (DSS) 40kD beads (Figure 4b). This result suggests that ABCB7 is required for protection against cytotoxic substrates like bortezomib and actinomycin D, (as well a vinblastine and paclitaxel, see Supplement Figure 4) rather than for proliferation. These results demonstrate that both CG32091 and ABCB7 are required for the multidrug resistance phenotype observed in esg+ stem/progenitor cells.

**Figure 4.**
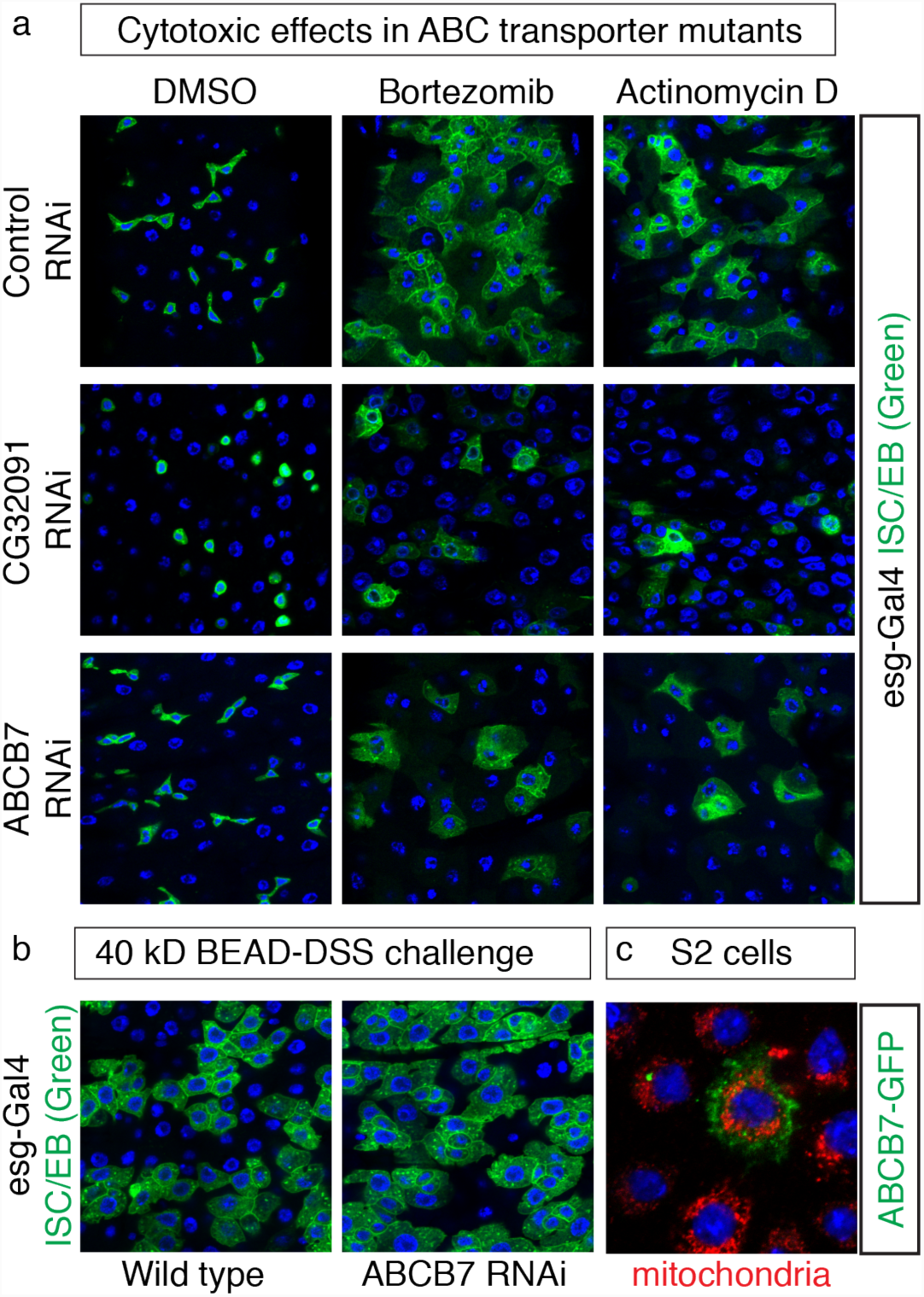
ABCB7 and CG32091 confer multidrug resistance to stem and progenitor cells. **(a)** Effect of RNAi knockdown of ABCB7 and CG32091 in stem/progenitor cells (green) challenged with cytotoxic drugs. **(b)** ABCB7 RNAi does not prevent stem/progenitor compensatory growth in the presence of the chemical EC-specific irritant DSS attached to 40kD dextran beads. **(c)** ABCB7-GFP localization (green) in transiently transfected S2 cells stained with the mitochondrial antibody Anti-ATP5A (red).

Our results are the first, to our knowledge to demonstrate that non-vertebrate stem cells are also multidrug resistant. This finding raises many questions about the specific ABC transporters that we identified, CG32091 and ABCB7, as well as general questions about the evolutionary pressures that keep stem cells but not their daughter cells primed for protection against exogenous cytotoxins. For example, while CG32091 and ABCB7 have both been identified in stress-response screens in Drosophila^12,38^, their mammalian homologs have not been implicated in stress or drug resistance. In fact, ABCB7 is highly conserved and required for heme transport across the inner mitochondrial membrane^39^. However, since ABC transporters tend to have multiple substrates and multiple roles, it is conceivable that the mammalian homologs of CG32091 and ABCB7 have drug resistance functions that will be detected in future assays, as has been for 19 of the 49 transporters encoded in the human genome^40^. In fact, we found that an ABCB7-GFP fusion protein localizes to the cell periphery and not to mitochondria in Drosophila S2 cells (Figure 4c), raising the possibility that it may localize differently in different cell types, as shown for the related “mitochondrial” transporter ABCB6^41^. Collectively, our results show that stem cells but not their differentiated daughter cells are under pressure to be able to express high levels of ABC transporters under cytotoxic conditions. While the specific ABC transporters may be different between different cell types and species, the net effect of having differentiated daughter cells remain vulnerable while stem cells protect themselves appears to be a general trend, whose mechanisms, costs, and benefits we anticipate will be dissected with the tools of Drosophila genetics.

## Acknowledgements

We thank Kim Tremblay for critically reading the manuscript. HD, JD, and ED received support from UMass undergraduate Commonwealth Honors College; ED is supported by a Goldwater Undergraduate scholarship.

## Author contributions

HD, JD, and MM conceived the project; HD designed the efflux assays; HD, JD, KK, OW, AB, KB, EH, MI, ED, VL, and RV performed experiments and analyzed results; SK and JD built the Gal4 knock in constructs; HD, JD, ED, and RV provided critical feedback on the manuscript; MM wrote the paper and supervised the project.

